# Laminin heparin-binding peptides promiscuously bind growth factors and enhance diabetic wound healing

**DOI:** 10.1101/259481

**Authors:** Jun Ishihara, Ako Ishihara, Kazuto Fukunaga, Priscilla S. Briquez, Jeffrey A. Hubbell

## Abstract

Laminin, as a key component of the basement membrane extracellular matrix (ECM), regulates tissue morphogenesis. We show that multiple laminin isoforms promiscuously bind to growth factors (GFs) with high affinity, through their heparin binding domains (HBDs) located in the a chain LG domains. Interestingly, these domains also bind to syndecan cell-surface receptors, promoting attachment of fibroblasts and endothelial cells. We next explore application of these multifunctional laminin HBDs in skin healing in the type 2 diabetic mouse. We demonstrate that covalent incorporation of laminin HBDs into fibrin matrix enables the slow-release of GFs. Incorporation of the α3_3043-3067_ laminin HBD significantly enhances *in vivo* wound-healing efficacy of vascular endothelial cell growth factor (VEGF)-A165 and platelet-derived growth factor (PDGF)-BB, under conditions where the GFs alone in fibrin are inefficacious. This laminin HBD peptide may be clinically useful by improving biomaterials as both GF reservoirs and cell scaffolds, leading to effective tissue regeneration.

## Introduction

Laminins are the most abundant glycoproteins of the basement membrane extracellular matrix (ECM) and can be found in almost all tissues of the body. They play essential roles in establishment of tissue architecture and stability, and provide cells with a structural scaffold. As such, laminins are involved in a variety of biological processes ranging from tissue survival, angiogenesis^1^ and neural development^2^, to skin re-epithelialization and wound healing^3–5^, and even cancer metastasis^6,7^. Laminins have been shown to regulate core cellular activities, such as adhesion, apoptosis, proliferation, migration and differentiation. Laminin is structured as a heterotrimer comprising three chains, a, b, and g, that assemble into a cross shape^3,8,9^. At least 16 different isoforms of laminin exist, made by various combinations of the five a (LAMA1–5), three b (LAMB1–3), and three g (LAMC1–3) chains that have been identified. Isoforms are accordingly named by their -αβγ chains: for instance, laminin-332 contains α3, β3 and γ2 chains. The differential expression of laminin isoforms depends on tissue type and state^1,10^.

In skin, for example, the epithelial basement membranes contain laminin-111 and laminin-211 during embryogenesis, but predominantly laminin-332 and laminin-511 in adults, the latter isoform expression seeming to diminish with age. Interestingly, dermal fibroblasts can transiently re-express laminin-211 after wounding^11^. Moreover, dermal blood vessels specifically express laminin-511 and laminin-311, in addition to the laminin-411 commonly found in endothelial basement membrane^3^.

In skin wound healing, laminin has been shown to have a critical role in re-epithelialization and angiogenesis^1,3^. Indeed, laminin a chain possesses 5 laminin-type G domain (LG) modules at the C-terminus, arranged in a tandem array, that are differentially processed under homeostatic conditions or during tissue repair^8^. In fact, laminin α3, α4, and α5 chains are physiologically cleaved by proteases, such as plasmin and elastase, in the linker sequence between LG3 and LG4 domains^12–15^, and processing in this region has been shown to correlate with the speed of wound closure^3^. As an example, laminin-332 is present in a cleaved form under homeostatic conditions; however, the expression of laminin-332 is upregulated after injury and the LG4 and LG5 domains are subsequently more present in wounds^1,12^. The release of LG4-LG5 module has been demonstrated to undergo further processing that releases bioactive peptides^12^, which promote blood vessel formation and keratinocyte migration, notably via syndecan binding^1,16–18^. In addition, laminin LG modules have been shown to bind to heparin sulfate, perlecan and fibulin-1^19^, as well as to a number of integrins, e.g. α3β1, α6β1, α7β1 and α6β4^20^.

In the past, ECM glycoproteins, including laminin, have been mainly considered for their biomechanical role in providing substrates for cell adhesion and migration, via direct interactions with cell-surface receptors^21^. Later, some ECM proteins have raised interest for their ability to regulate the partitioning and bioavailability of soluble signaling molecules within tissues, thus highlighting a new role for the ECM in coordinating the spatio-temporal release of these molecules. Examples of such soluble signals are growth factors, which are key morphogenetic proteins broadly involved in the control of core cellular behaviors, and which have been shown to be crucial for wound healing^22–24^. Particularly, fibronectin^25,26^, vitronectin^27^, and fibrinogen^28^ have been reported to directly bind to GFs, which control their release kinetics *in vivo,* acting as a GF reservoir. Importantly, it has been noticed that GF binding occurs at the heparin-binding domains (HBDs) of ECM glycoproteins, especially in fibronectin^1,26^, vitronectin^27^, fibrinogen^28^ and osteopontin^29^. Sustained release of GFs from the ECM eventually enhances and prolongs GF-receptor signaling. Furthermore, the proximity of GF-binding sites and integrin-binding sites in some ECM glycoprotein chains, as in fibronectin or vitronectin, can induce synergistic GF signaling by clustering GF receptors and integrin at the cell surface.

Despite the importance of laminin in the ECM composition and its role in tissue morphogenesis, interactions between laminin and GFs have been poorly investigated to date. Therefore, in this study, we aim to elucidate direct interactions between laminin and GFs; we hypothesized that laminin could bind to several GFs, as observed in other ECM glycoproteins, and that these interactions could take place at laminin HBDs, some of which having been already identified in LG domains^30–32^. We then sought to demonstrate the GF reservoir role of laminin *in vivo*, and its potential to control the delivery of GF from fibrin materials in chronic wound healing, as a clinically-relevant model of tissue repair.

## Results

### Multiple GFs bind to multiple isoforms of laminin

We first examined the capacity of a variety of full-length laminin isoforms (−111, −211, −332, −411, −421, −511, and −521) to bind GFs from the VEGF/PDGF, FGF, BMP, NT, IGF, EGF and CXCL chemokine families. Binding of laminin to absorbed GFs was detected using an antibody against laminin, and signals greater than 0.1 were considered to be indicative of a binding event. Overall, we found that multiple GFs strongly bound to all tested laminin isoforms (Fig. 1A). Specifically, from the VEGF/PDGF family, VEGF-A165, PlGF-2, PDGF-AA, PDGF-BB, and PDGF-CC bound to all isoforms of laminin, in contrast to VEGF-A121, PlGF-1, and PDGF-DD which did not show binding. From the FGF family, we observed that FGF-2, FGF-7, FGF-10, and FGF-18 bound to all laminin isoforms, whereas FGF-1, FGF-6, and FGF-9 did not. Among the BMPs, BMP-2 and BMP-3 showed binding to laminins, but not BMP-4 and BMP-7. NT-3 and BDNF showed strong binding towards all tested laminin isoforms, while bNGF bound only weakly. Neither IGF-1 nor IGF-2 displayed significant binding to laminins. In addition, HB-EGF weakly bound to laminins. As to the tested chemokines, CXCL-12γ bound to all laminin isoforms, whereas CXCL-11 and CXCL-12α bound weakly to laminin-332 but not to the other isoforms.

**Fig. 1.**
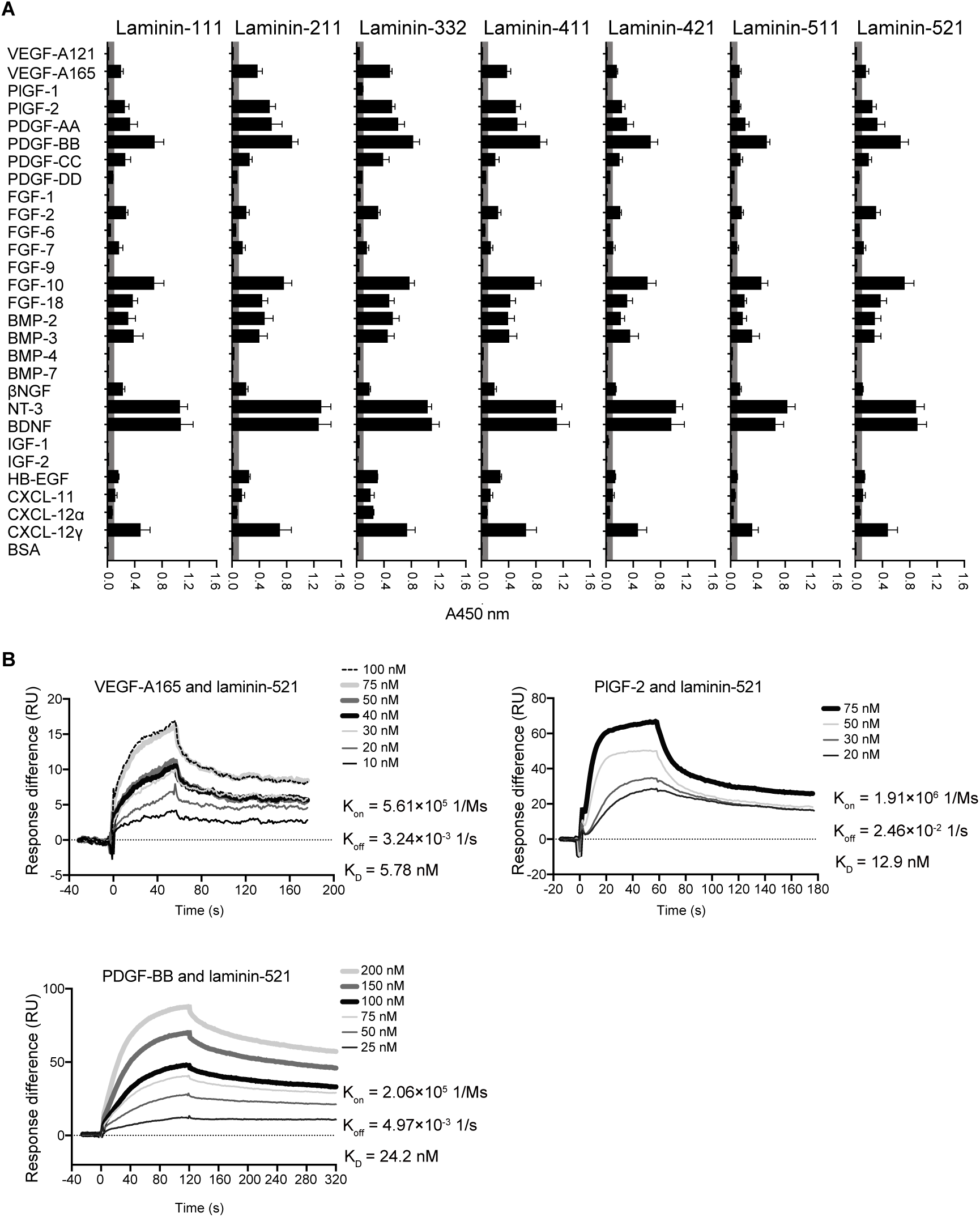
Multiple isoforms of laminin bind promiscuously to GFs and chemokines with high affinities. (A) Multiple isoforms of full-length laminin (−111, −211, −332, −411, −421, −511, and −521) binding to GFs and CXCL chemokines were measured by ELISA. A450 nm represents absorbance at 450 nm. BSA-coated wells served as negative controls (n = 4, mean ± SEM). Signals greater than 0.1 (grey box) are considered to be significant. (B) Affinities (K_D_ values are shown) of full-length laminin against VEGF-A165, PlGF-2 and PDGF-BB were measured by SPR. A SPR chip was functionalized with laminin-521 (~2000 RU), and each GF was flowed over the chip at indicated concentrations. Curves represent the specific responses (in RU) to laminin obtained. Experimental curves were fitted with Langmuir binding kinetics. Binding kinetics values [dissociation constants (K_D_) and rate constants (K_on_ and K_off_)] determined from the fitted curves are shown.

Next, we measured the affinities between laminin-521, as an example, and VEGF-A165, PlGF-2, and PDGF-BB using surface plasmon resonance (SPR). SPR chips were functionalized with laminin-521, and growth factors were flowed over the surface. The obtained binding curves were fitted with Langmuir binding kinetics to calculate specific dissociation constants (K_D_) (Fig. 1B). K_D_ values were 5.8 nM for VEGF-A165, 12.9 nM for PlGF-2, and 24.2 nM for PDGF-BB. The nM range of K_D_ values demonstrated the strong binding affinities of laminin-521 to the selected GFs.

### GFs bind to the HBDs of laminin

Because the GFs that bound to laminins have also been previously reported to bind to other ECM glycoproteins through HBDs^25,28,33^, we hypothesized that HBDs of laminins might be responsible for the interactions between GFs and laminin. To address this hypothesis, ELISA assays were repeated for VEGF-A165, PlGF-2 or FGF-2 in the presence of heparin added in excess (10 µM). As a result, we observed that heparin inhibited GFs binding to laminin (Fig. 2A–C), supporting that laminin HBDs mediated interactions with GFs. To further confirm this, we tested direct GF binding to the LG domains from human laminin α3, α4 and α5, within which HBDs of laminin were localized^30^. We found that VEGF-A165, PlGF-2, PDGF-BB, and FGF-2 bound to laminin LG domains α3_2928-3150_, α4_826-1816_ and α5_3026-3482_, in contrast to VEGF-A121 and PlGF-1 which did not show any binding (Fig. 3A–C), as tested by ELISA. The binding affinities between α3_2928-3150_ and VEGF-A165 or PDGF-BB were then measured by SPR, and K_D_ values were 1.2 nM for VEGF-A165, and 10.2 nM for PDGF-BB (Fig. 3D). These data again demonstrated the strong affinities of the laminin LG domain to the tested GFs.

**Fig. 2.**
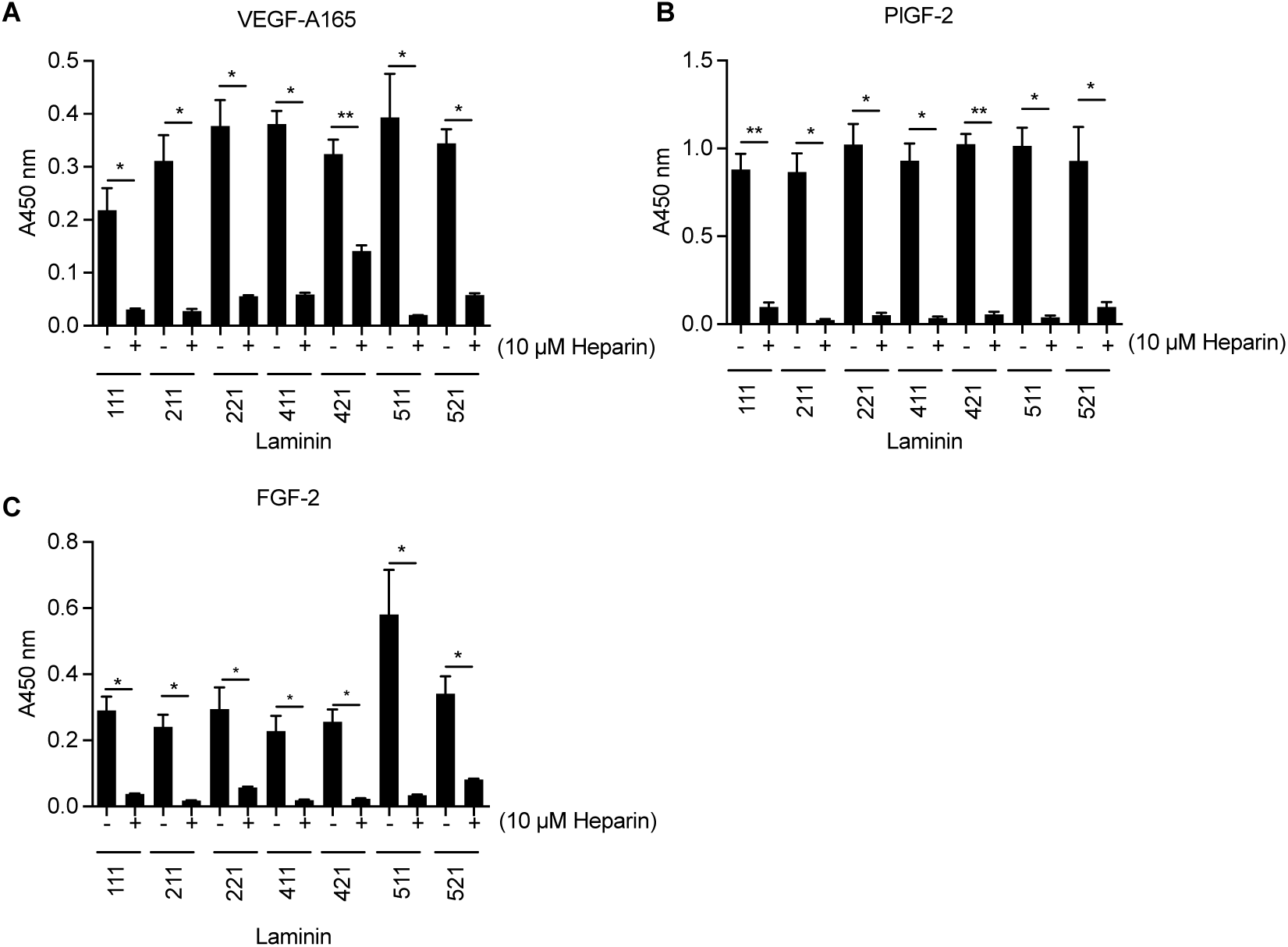
Heparin inhibits GF-laminin binding. Inhibition of GF-binding to laminin (−111, −211, −221, −411, −421, −511, and −521) by excess heparin. ELISA plates were coated with 10 µg/mL laminin and further incubated with a 1 μg/mL (A) VEGF-A165, (B) PlGF-2, or (C) FGF-2 solution in the absence or presence of excess (10 μM) heparin. Bound GFs were detected using a specific antibody for each GF (n = 4, mean ± SEM). Statistical analyses were done using Mann-Whitney U test by comparing the signals with and without heparin. *p < 0.05, **p < 0.01.

**Fig. 3.**
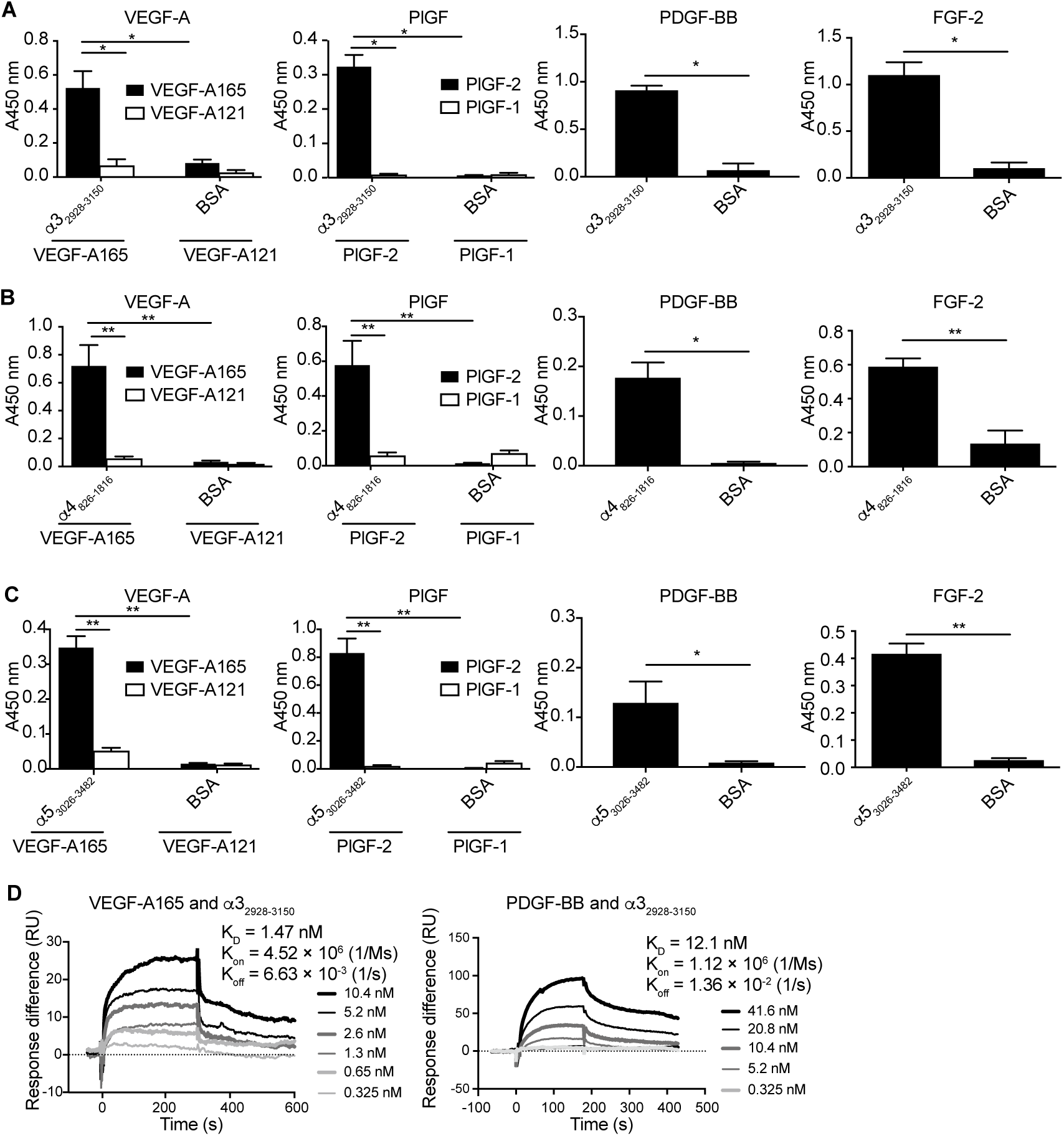
GFs bind to recombinant LG domain protein derived from laminin α3, α4 and α5 chains. Affinity of GFs against recombinant laminin LG domains. ELISA plates were coated with 1 µg/mL (A) α3_2928-3150_, (B) α4_826-1816_, or (C) α5_3026-3482_ and further incubated with 1 μg/mL of VEGF-A165, VEGF-A121, PlGF-2, PlGF-1, PDGF-BB, or FGF-2 solution. Bound GFs were detected using a specific antibody for each GF (n = 4, mean ± SEM). Statistical analyses were done using the Mann-Whitney U test by comparing the signals obtained from the laminin domain- and the BSA- coated wells. *p < 0.05, **p < 0.01. (D) Affinities (K_D_ values are shown) of laminin α3_2928-3150_ against VEGF-A165 and PDGF-BB were measured by SPR. A SPR chip was functionalized with the laminin α3_2928-3150_ recombinant protein (~1000 RU), and each GF was flowed over the chip at indicated concentrations. Curves represent the specific responses (in RU) to laminin. Experimental curves were fitted with Langmuir binding kinetics. Binding kinetics values [dissociation constants (K_D_) and rate constants (K_on_ and K_off_)] determined from the fitted curves are shown.

We next examined the binding of GFs to chemically synthesized laminin LG domain peptides, the sequences of which are all derived from human laminin sequences (Table 1, Fig. 4A). These peptides are putative HBDs; they were determined based on previous reports with mouse or human HBD sequences^30^, or are positively charged sequences located within the linker domain between the LG3 and LG4 domains in laminin α3, α4 and α5 chains. Of 9 tested peptides, 6 bound to heparin (i.e. HBDs), namely α3_2932-2951_, α3_3043-3067_, α4_1408-1434_, α41521-1543, α53300-3330, and α53417-3436 among which α32932-2951, α41408-1434, and α5_3300-3330_ are derived from the LG3-LG4 linker. Interestingly, α5_3312-3325_, which is a subdomain of α5_3300-3330_, did not bind to heparin.

**Table 1.**
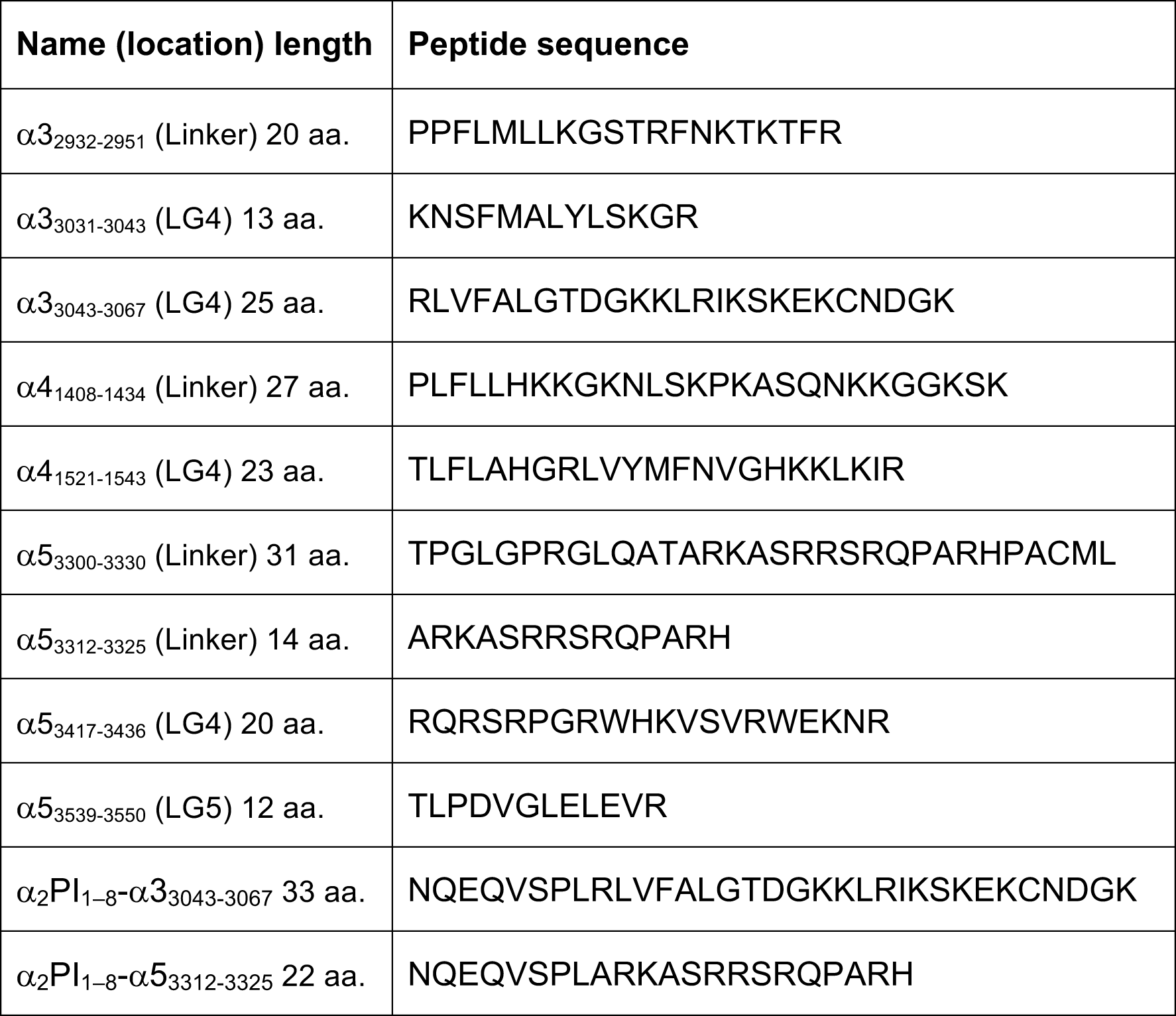
The sequences of laminin-derived peptides.

**Table 2.** Summary of laminin-derived peptides interactions.

| Laminin-derived peptides | Interaction with |  |  | Cell adhesion |  |
| --- | --- | --- | --- | --- | --- |
|  | Heparin | GFs | Syndecans | Fibroblasts | HUVECs |
| $\alpha 3_{2932-2951}$ | ++ | + | + | + | + |
| $\alpha 3_{3031-3043}$ | | + | + | + | |
| **$\alpha 3_{3043-3067}$** | ++ | ++ | ++ | ++ | ++ |
| $\alpha 4_{1408-1434}$ | ++ | ++ | ++ | | |
| $\alpha 4_{1521-1543}$ | ++ | + | ++ | + | + |
| $\alpha 5_{3300-3330}$ | ++ | + | ++ | | |
| $\alpha 5_{3312-3325}$ | | | + | | |
| $\alpha 5_{3417-3436}$ | ++ | ++ | ++ | + | |
| $\alpha 5_{3539-3550}$ | | + | | | |

++ indicates high affinities, + indicates medium/low affinities. The laminin-derived peptide tested *in vivo* is highlighted in gray.

**Fig. 4.**
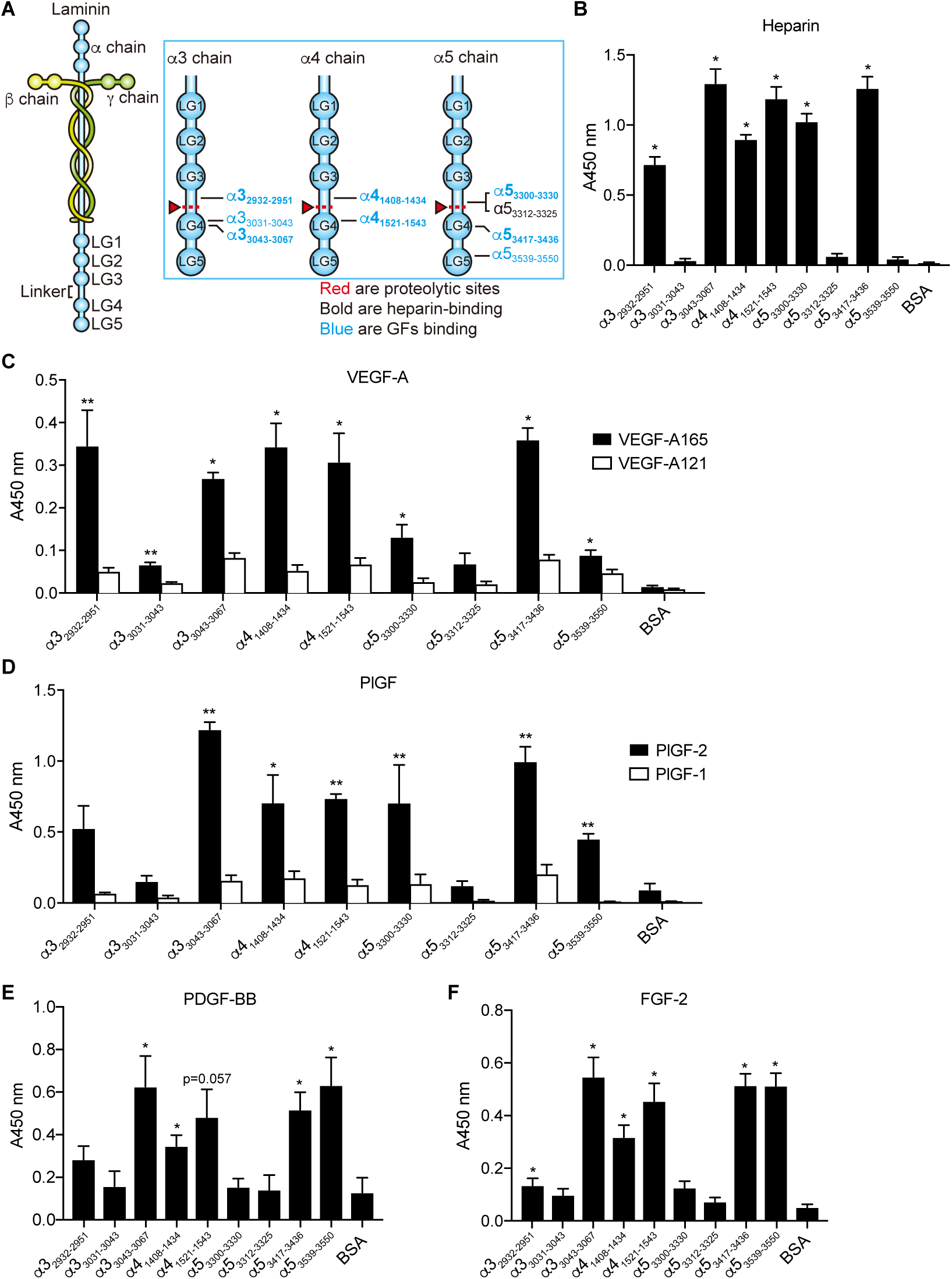
GFs bind to chemically synthesized laminin HBD peptides derived from the LG domain of laminin α3, α4, and α5 chains. (A) The location of laminin-derived peptides in the LG domain of laminin α3, α4, and α5 chains. (B-F) Affinity of heparin and GFs against chemically synthesized peptides derived from the LG domain of laminin α3, α4, and α5 chains. ELISA plates were coated with 10 µg/mL laminin peptide and further incubated with (B) biotinylated heparin, (C) VEGF-A165 and VEGF-A121, (D) PlGF-2 and PlGF-1, (E) PDGF-BB, or (F) FGF-2. Concentrations were 1 μg/mL for GFs and 10 µg/mL for heparin. Bound heparin was detected with streptavidin, and bound GFs with a specific antibody for each GF (n = 4, mean ± SEM). Statistical analyses were done using Mann-Whitney U test by comparing the signals obtained from the laminin peptide- and the BSA-coated wells. *p < 0.05, **p <0.01.

Finally, the affinities of VEGF-A, PlGF, PDGF-BB, and FGF-2 to these peptides were examined (Fig. 4B–F). We observed that all heparin-binding peptides showed significant binding to some GFs. Indeed, α3_3043-3067_, α4_1408-1434_, and α5_3417-3436_ bound to VEGF-A165, PlGF-2, PDGF-BB, and FGF-2. α4_1521-1543_ showed similar results except for the binding to PDGF-BB, which was not statistically significant. α3_2932-2951_ and α5_3300-3330_ preferentially bound to VEGF-A165 and FGF-2, and VEGF-A165 and PlGF-2 respectively. As to the non-heparin-binding peptides, α5_3312-3325_ did not show particular binding to any tested GF. Interestingly, α5_3539-3550_, which did not show binding to heparin, significantly bound to all tested GFs, and α3_3031-3043_ bound to VEGF-A165. None of the tested laminin-derived peptides bound to VEGF-A121 nor to PlGF-1, consistent with the results obtained in Fig. 1 and Fig. 3. Taken together, these data suggest that GFs bind to the HBDs of laminin, located in the LG3-LG4 linker or in LG4-LG5 domains.

### Laminin HBD peptides promote adhesion of multiple types of cells

Because the laminin HBDs have been reported to bind to syndecan^30^, a key cell surface adhesion molecule, we tested syndecan binding to the synthesized laminin-derived peptides (Fig. 5A–D). α3_3043-3067_, α4_1521-1543_, α4_1408-1434_, α5_3417-3436_, and α5_3300-3330_ showed significant binding to all isoforms of recombinant syndecans, i.e. syndecan 1–4. α3_2932-2951_, α3_3031-3043_, and α5_3312-3325_ showed weak binding to the tested syndecans, while α5_3539-3550_ did not show binding to any syndecan isoform. Because laminin-derived peptides that interact with syndecans may further promote cell adhesion by providing binding substrates, we tested fibroblasts and HUVEC adhesion to plates coated with these peptides. We indeed observed enhancement of fibroblast attachment on α3_2932-2951_, α3_3031-3043_, α3_3043-3067_, α4_1521-1543_ and α5_3417-3436_-coated surfaces (Fig. 6A). Fibroblast binding was observed even after EDTA-treatment, consistent with syndecan function (Fig. 6B). Of these peptides, α3_2932-2951_, α3_3043-3067_, and α4_1521-1543_ also promoted HUVEC attachment (Fig. 6C), even in presence of EDTA in the case of α3_3043-3067_ (Fig. 6D). Interestingly, peptides that promoted both fibroblast and HUVEC adhesion *in vitro* through syndecan binding were those that we previously found to be laminin HBDs (Fig 4A).

### Retention of VEGF-A165 and PDGF-BB in fibrin matrix is increased by the incorporation of laminin HBDs peptides

We then sought to determine whether laminin HBD peptides, which showed binding to GFs, were able to improve the retention of VEGF-A165 and PDGF-BB within fibrin matrix. These GFs are known to be quickly released from fibrin matrices upon delivery, which limits their wound healing efficacy *in vivo*^26,28,33^. For this purpose, we selected α3_3043-3067_ and α5_3417-3436_ laminin HBD peptides, and fused them to a transglutaminase-reactive sequence from the α_2_-plasmin inhibitor^26^ to allow their covalent incorporation by factor XIIIa into fibrin matrices during polymerization. GF release from fibrin matrices containing α_2_PI_1-8_-α3_3043-3067_, α_2_PI_1-8_-α5_3417-3436_ or no laminin-derived peptide were then monitored daily and quantified by ELISA (Fig. 7A, B). As expected, we observed that VEGF-A165 and PDGF-BB were quickly released from the fibrin matrix (> 85% released after 24 h). However, incorporation of either α_2_PI_1-8_-α3_3043-3067_ or α_2_PI_1-8_-α5_3417-3436_ allowed significant retention of VEGF-A165 and PDGF-BB into matrices, which were respectively released at VEGF-A165 (α_2_PI_1-8_-α3_3043-3067_: 25%, α_2_PI_1-8_-α5_3417-3436_: 31%) and PDGF-BB (α_2_PI_1-8_-α3_3043-3067_: 45%, α_2_PI_1-8_-α5_3417-3436_: 47%) after 5 days. This data highlights the key biological role of laminin in sequestering GFs into ECM, and demonstrates the potential of laminin HBD peptides to control GF delivery from fibrin biomaterials (Fig. 7A, B).

### Laminin HBD-functionalized fibrin matrices potentiate GFs and promote wound healing *in vivo*

We further evaluated whether fibrin matrices engineered with laminin-HBD peptides could enhance skin repair in a model of delayed wound healing, by controlling the release of VEGF-A165 and PDGF-BB *in vivo*. More precisely, VEGF-A165 (100 ng/wound) and PDGF-BB (50 ng/wound) were co-delivered from fibrin matrix onto full-thickness back-skin wounds in type 2 diabetic db/db mice, which provides a well-established and clinically-relevant model of impaired wound healing^26^. Here, we particularly functionalized fibrin with the laminin peptide α3_3043-3067_, since it bound to GFs and syndecans, and promoted fibroblast and endothelial cells adhesion *in vitro* (Fig. 4–6). Four groups were tested: fibrin only, fibrin functionalized with α_2_PI_1–8_-α3_3043-3067_, fibrin containing GFs, and fibrin functionalized with α_2_PI_1–8_-α3_3043-3067_ and containing GFs. Wound histology was analyzed after 7 days, considering that wounds are normally fully closed after 15 days when treated with fibrin matrix^26^. As a result, wounds that received fibrin matrices containing GFs or α_2_PI_1–8_-α3_3043-3067_ peptide only did not differ from wounds treated with fibrin alone, neither in amount of granulation tissue nor in extent of wound closure (the latter indicated by re-epithelialization) (Fig. 7C, D). In contrast, the co-delivery of VEGF-A165 and PDGF-BB in fibrin functionalized with α_2_PI_1–8_-α3_3043-3067_ led to a significantly faster wound closure after 7 days, as well as a significant increase in granulation tissue formation (Fig. 7C, D). Representative wound morphology for all four treatments is presented in Fig. 7E. Clear differences in granulation tissue thickness and extent of re-epithelialization can be visualized when GFs were delivered within the α_2_PI_1–8_-α3_3043-3067_ peptide-functionalized fibrin matrix compared to the other conditions. This set of data indicates that α_2_PI_1– 8_-α3_3043-3067_ improved the GF delivery capacity of fibrin *in vivo*, resulting in an enhanced wound healing.

## Discussion

As a cell scaffold protein, laminin tightly regulates cell adhesion, motility, survival and differentiation, thus playing a critical role in tissue homeostasis and wound healing ^8^. In fact, although the expression of particular laminin isoforms depends on the tissue type and state, laminins reportedly promote tissue repair in muscle, nerve, liver, and skin^34^. In this study, we uncovered a novel property of laminin, showing that multiple laminin isoforms promiscuously bind to heparin-binding GFs with high affinities (Fig. 1). Interactions between GFs and the ECM are known to be essential for controlling GFs release kinetics *in vivo*, which strongly modulates tissue morphogenesis^26^. While sequestration of GFs within basement membranes was previously reported to be mediated through binding to glycosaminoglycans^35^, our data demonstrates that laminin can serve this GF reservoir function as well.

Interestingly, most of the GFs that bind to laminin have also been shown to bind to other ECM proteins, such as fibrinogen and fibronectin, suggesting similar binding mechanisms^25,26,28,33^. However, a few exceptions can be noticed; for example, PDGF-CC showed specific binding to laminin, but neither to fibrinogen nor fibronectin^28,33^. In contrast, BMP-7, which binds specifically to fibronectin^25^ but not fibrinogen^28^, did not show binding to laminin. Moreover, kinetic analysis of laminin isoform-521 binding towards VEGF-A165, PDGF-BB and PlGF-2 revealed K_D_ in nM range, highlighting the broad yet high affinity interactions between laminin and GFs.

Furthermore, we identified GF binding to laminin to mainly occur at heparin-binding sites, by showing that heparin directly compete with GF-laminin interactions and dramatically reduced them when added at high concentration (Fig. 2). Additionally, all laminin HBDs tested in this study were able to bind to GFs (Fig. 3, 4). Yet, unexpectedly, GFs binding to laminin does not seem to be limited to HBDs, as a few non-heparin binding peptides also bound to some GFs, notably α3_3031-3043_ and α5_3539-3550_. These peptides are human alignments of reported mouse HBD peptides, called A3G75 and A5G94 respectively^30^. Although they do not show heparin binding under our experimental conditions, they may bind to GFs via another mechanism. Thus, the mechanism of GF-binding to laminin still remains incompletely clarified, and may be resolved by further crystallography studies of GF-laminin complex.

Physiologically, proteolytic cleavage of LG4 and LG5 domains is crucial for the deposition of laminin in the native ECM^12,14,36^. Upon tissue injury, laminin is overexpressed, and LG4-LG5 domains accumulate in wounds^1,12^, wherein they promote tissue healing mechanisms^37^. In this study, we particularly characterized laminin-derived peptides that are located just before the proteolytic cleavage site, in the linker between the LG3 and LG4 domains, or within the LG4-LG5 domains (Table. 2, Fig. 4A). On one side, we discovered 3 novel heparin-, GF- and syndecan-binding peptides within the LG3-LG4 linker regions of α3, α4, and α5 chains, namely α3_2932-2951_, α4_1408-1434_, and α5_3300-3330_, identifiable through their highly cationic sequences (Fig. 4). Since α3, α4 and α5 chains are known to be predominantly present in their processed form (i.e. lacking LG4-LG5) in mature, unwounded skin^3,12–15^, it is likely that these peptides are exposed *in vivo* under homeostatic conditions, thus providing both GF ligands and cell adhesion sites in basement membranes. Interestingly, laminin α1 chain, which is not proteolytically processed^13^, and α2 chain do not contain such cationic sequences in the LG3-LG4 linker region, which might reflect functional differences between α chain isoforms. On the other side, we identified 5 peptides in the LG4 and LG5 domains of α3, α4 and α5 chains that displayed specific binding to GFs, in particular to VEGF-A165. Among them, α3_3043-3067_, α5_3539-3550_, and α5_3417-3436_ additionally bound to PDGF-BB, FGF-2 and PlGF-2 with high affinities (Fig. 4). These growth factors are well-known as key regulators of the wound healing cascade, and are particularly involved in wound angiogenesis. Therefore, we propose that the reported positive effects of LG4-LG5 domains during wound healing might be related to promiscuous interactions with GFs, in addition to via binding to syndecans and release of laminin-derived pro-angiogenic peptides^1,16,17^.

Previous studies have shown that the formation of ECM protein:GF complexes can synergistically enhance GF receptor signaling^26,27^. For example, simultaneous presentation of GFs and integrin-binding sites by an engineered fibronectin fragment (namely FN III9-10/12–14) incorporated into fibrin drastically improved the effect of VEGF-A165 and PDGF-BB on skin repair^26^. Here, we identified 5 laminin HBDs that are able to bind to both GFs and syndecan cell-receptors (Fig. 4 and 5), among which α3_3043-3067_, α4_1521-1543_ and α5_3417-3446_ further promoted cell attachment (Fig. 6). Although syndecans are not known to directly activate major signaling pathways, they support cell adhesion and integrin signaling^18^. Moreover, direct binding of laminin peptides from LG domains to integrins has also been reported; for example, the integrin α3β1 binds to α3_2932-2943_^38^. Nevertheless, in our assays, EDTA did not abolish cell adhesion, suggesting that initial cell attachment was mediated by syndecans rather than integrins (the binding of which is Ca^2+^-dependent). Consequently, and considering the short length of the laminin HBD peptides, it is unlikely that laminin HBD peptides can enhance GF signaling via synergy with integrins. However, we believe that both GF- and cell-binding properties of laminin HBDs substantially contribute to the promotion of wound healing, and so propose a potential application for laminin HBDs in the treatment of chronic diabetic ulcers (Fig. 7).

**Fig. 5.**
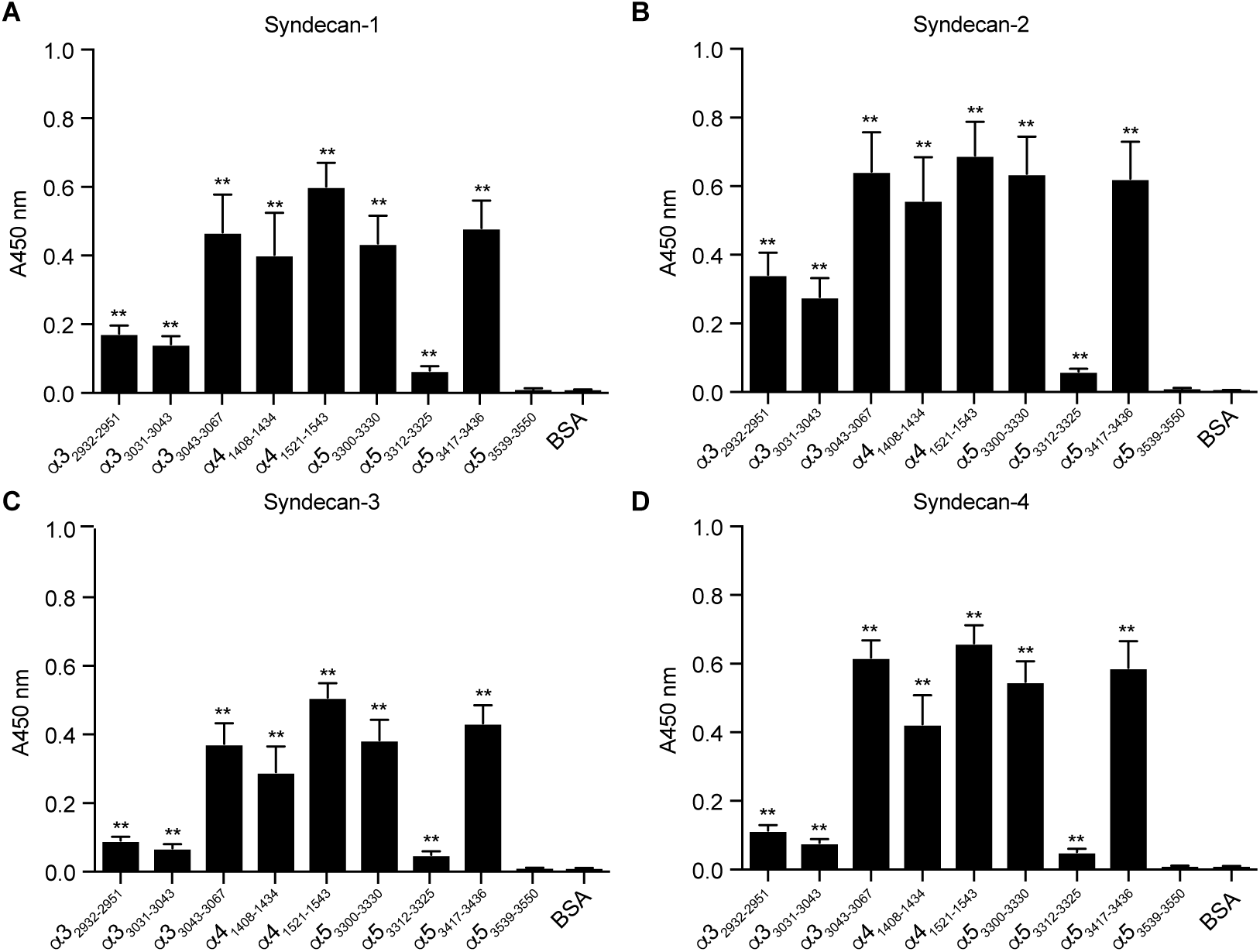
Chemically synthesized peptides derived from the LG domain of laminin α3, α4 and α5 chains bind to syndecans. Affinity of syndecans to chemically synthesized peptides derived from the laminin α3, α4 and α5 LG domains. ELISA plates were coated with 10 µg/mL laminin peptide and further incubated with 1 μg/mL of (A) syndecan-1, (B) syndecan-2, (C) syndecan-3, or (D) syndecan-4. Bound syndecans were detected using an antibody against histidine-tag on the recombinant syndecans (n = 8, mean ± SEM). Statistical analyses were done using Mann-Whitney U test by comparing the signals obtained from the laminin peptide- and the BSA-coated wells. *p < 0.05, **p < 0.01.

**Fig. 6.**
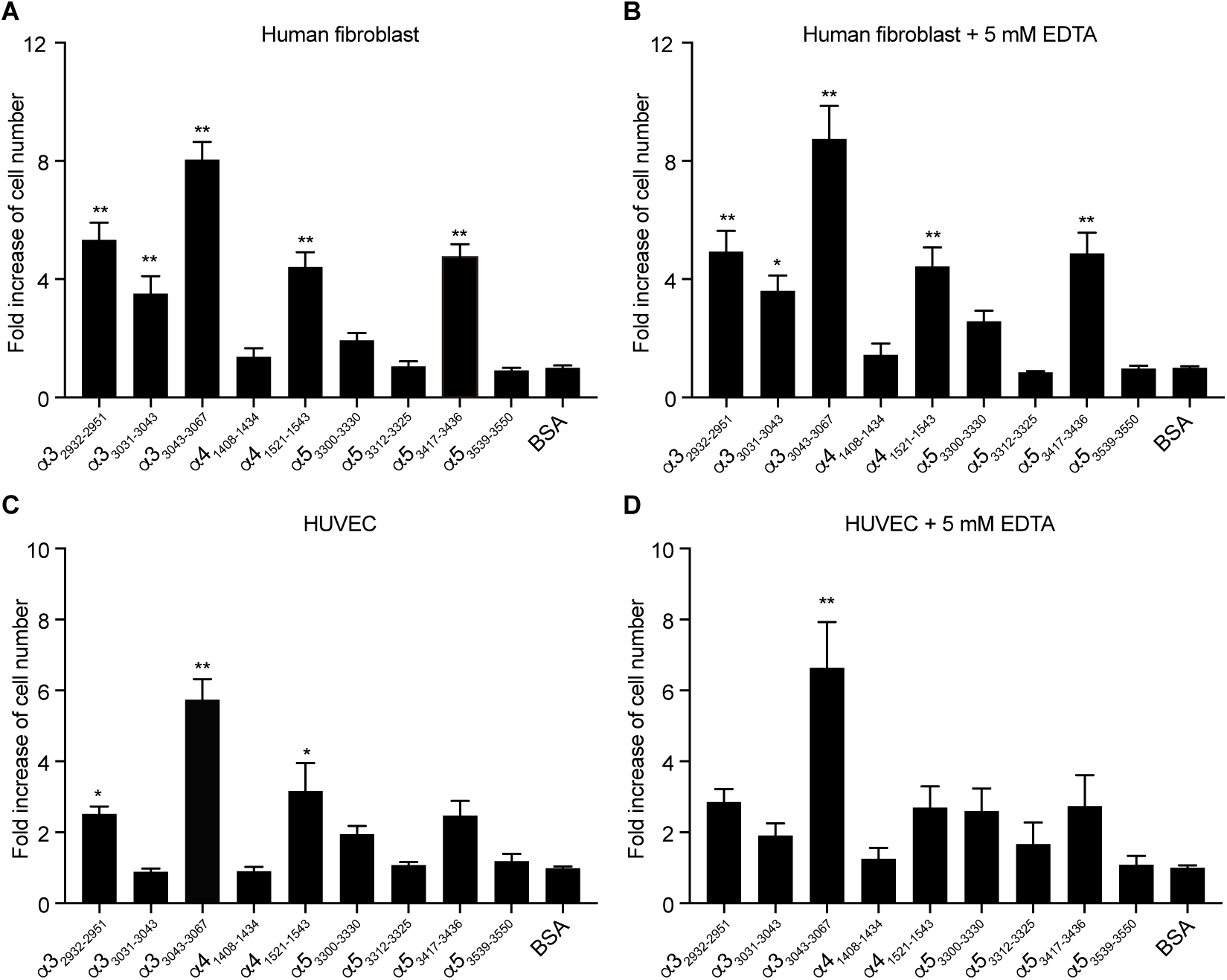
Laminin HBD peptides promote fibroblast and endothelial cell adhesion *in vitro*. A, B) 3000 cells/well human lung fibroblasts were cultured (A) without or (B) with 5 mM EDTA in FGM-2 culture media containing 1% FBS. (C, D) 3000 cells/well HUVEC were cultured (C) without or (D) with 5 mM EDTA in EBM-2 culture media containing 100 ng/ml VEGF-A165 and 1% FBS. Cells were plated on 1 μg/mL laminin peptide pre-coated non-tissue culture treated plates and incubated for 30 min at 37°C. After plate washes, cell numbers were quantified using a CyQUANT assay (n = 10, mean ± SEM). The signals obtained from BSA-coated wells are normalized to 1, and relative fold increases of cell numbers were calculated. Statistical analyses were done using ANOVA with Tukey’s test. Kruskal-Wallis test followed by Dunn’s multiple comparison was used in (B, C). *p < 0.05, **p < 0.01.

**Fig. 7.**
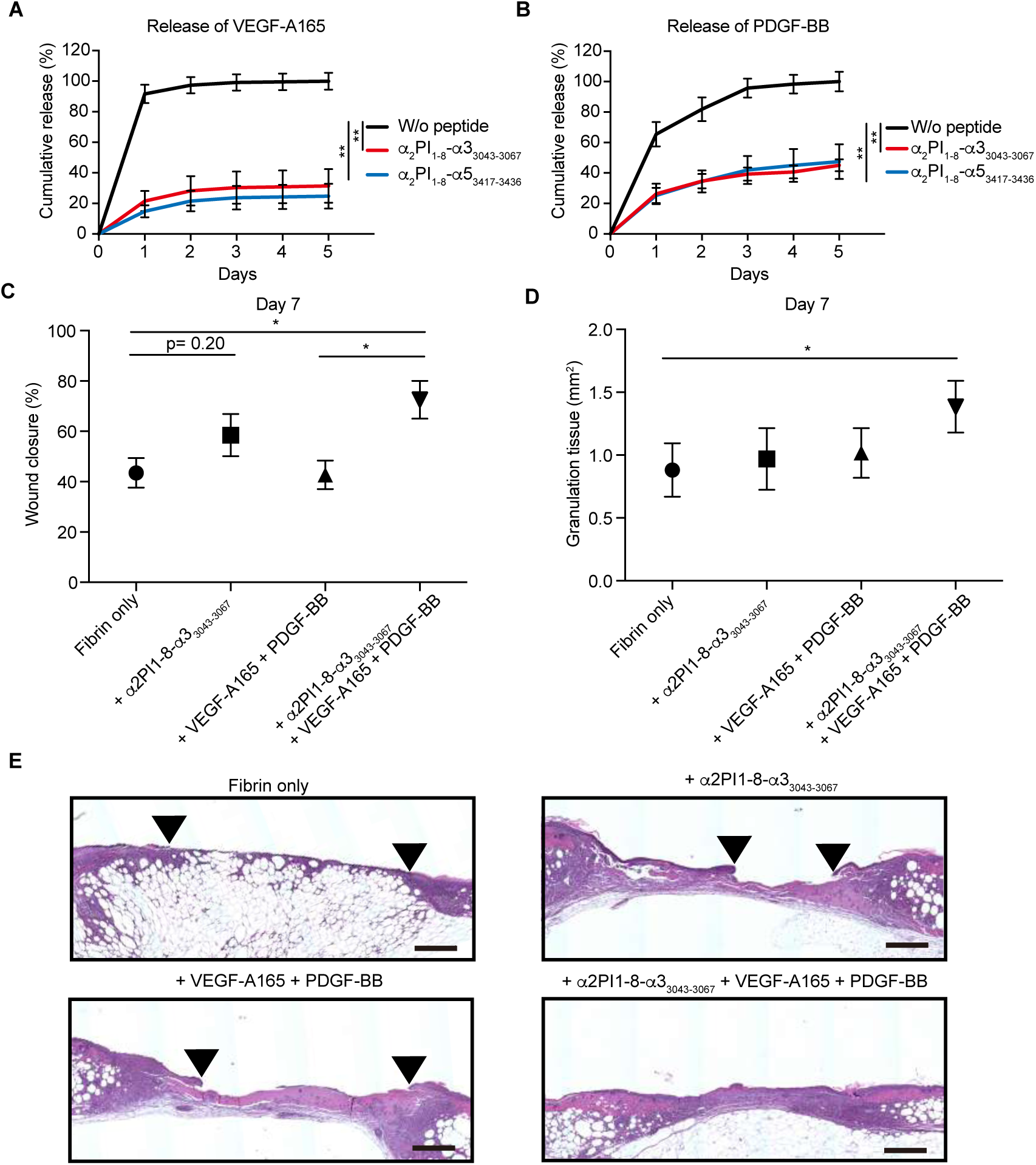
Delivering GFs within laminin HBD peptide-functionalized fibrin matrices enhances skin wound healing in db/db diabetic mice. (A,B) GF retention in fibrin matrix. α_2_PI_1–8_-α3_3043-3067_ or α_2_PI_1–8_-α5_3417-3436_ peptide-functionalized fibrin matrices were made in the presence of VEGF-A165 or PDGF-BB, and further incubated in 8 volumes of physiological buffer for 5 days. The buffer was changed each day, and released GFs were quantified daily. Graphs show the cumulative release of (A) VEGF-A165 or (B) PDGF-BB over 5 days (n = 4; mean ± SEM). All data points for laminin HBD peptides were statistically significant compared to controls without laminin HBD peptide (p < 0.01, Mann-Whitney U test). (C-F) Full-thickness back-skin wounds were treated with combined VEGF-A165 (100 ng/wound) and PDGF-BB (50 ng/wound). Four groups were tested: fibrin only, fibrin functionalized with α_2_PI_1–8_-α3_3043-3067_ peptide, fibrin containing admixed GFs, and fibrin functionalized with α_2_PI_1– 8_-α3_3043-3067_ peptide containing GFs. After 7 days, (C) wound closure and (D) granulation tissue area were evaluated by histology (means ± SEM, fibrin only, n=10; other treatment groups, n = 11). *P < 0.05, ANOVA with Tukey’s test. (E) Wound histology (hematoxylin and eosin staining) at day 7. Red arrows indicate tips of epithelium tongue. The granulation tissue (pink-violet) is characterized by a large number of granulocytes with nuclei that stain in dark-violet or black. Muscle under the wounds is stained in red. Fat tissue appears as transparent bubbles. Scale bar = 800 µm.

Although GFs are promising drugs for tissue regeneration, their uncontrolled delivery upon application on wounded tissue had limited their clinical efficacy and safety to date^39,40^. For example, recombinant human VEGF-A has not been approved for clinical use by the U.S. Food and Drug Administration (FDA) due to a negative result in phase II clinical trials^41^. PDGF-BB (Regranex in the clinic) has shown clinical efficacy, but safety issues such as cancer risk have been flagged, potentially due to high dosing^1,42^. Thus, controlling GF delivery to improved efficacy and dose reduction seems essential in future GF-based therapies and could be achieved by use of biomaterials^26^. Here, we showed that covalent incorporation of an engineered GF-binding domain derived from laminin, α_2_PI_1–8_-α3_3043-3067_, into fibrin matrix significantly enhanced the effect of VEGF-A165 and PDGF-BB on skin wound healing, by highly increasing GF retention into fibrin (Fig. 7). In contrast, wounds treated with fibrin matrix containing GFs only, in which PDGF-BB and VEGF-A165 were not specifically retained in the fibrin matrices, had no detectable effect on wound healing at the tested dose (Fig. 7). We have previously shown that HBDs derived from fibrinogen (Fg b15-66_(2)_) and fibronectin (FN III9-10/12–14) promote wound healing in combination with GFs when incorporated into fibrin matrix^26,28^. A main advantage of using the laminin HBD peptide for this purpose, compared to the previously described GF-binding domains, is production simplicity: the laminin HBD peptide is short enough to be chemically synthesized in large scale, rather than requiring recombinant expression. Furthermore, we showed that a laminin HBD can functionalize fibrin matrix in both aspects as a GF reservoir and an adhesion-promoting cell scaffold (Fig. 6 and 7).

In conclusion, we found that multiple isoforms of laminin promiscuously bind to a number of GFs from the VEGF/PDGF, FGF, BMP, and NT families, in addition to HB-EGF and CXCL12γ, through their HBDs. By engineering a fibrin matrix displaying the α3_3043-3067_ laminin HBD, as a proof example, we have demonstrated that the laminin HBD peptide promotes skin wound closure in the db/db mouse, as a model of delayed wound healing, when applied with VEGF-A165 and PDGF-BB. In addition to highlighting a GF-modulating function for laminin, an important tissue repair protein, we show that both GF- and cell-binding character promotes tissue repair when incorporated within fibrin matrix, which may be clinically useful.

## Acknowledgements

We thank Prof. Y. Kikkawa and Prof. M. Nomizu (Tokyo University of Pharmacy and Life Sciences) for useful discussion, the Human Tissue Resource Center of the University of Chicago for histology analysis, and the Biophysics Core Facility of the University of Chicago for SPR analysis.

## Funding

This work was funded in part by the Chicago Biomedical Consortium with support from the Searle Funds at The Chicago Community Trust and by the US National Institutes of Health grant DP3DK108215 (to J.A.H.).

## Author contributions

J.I., and J.A.H. designed the project. J.I., A.I., and K.F. performed the experiments. J.I., A.I., K.F. and J.A.H. analyzed the data. J.I., P.S.B. and J.A.H. wrote the paper.

## Competing interests

The University of Chicago has filed for patent protection on some of the technology described herein, and J.I., A.I., P.S.B. and J.A.H. are named as inventors.

## Materials and methods

### Growth factors and chemokines

All growth factors (GFs) and chemokines were purchased in their mature forms, highly pure (> 95% pure), carrier-free, and lyophilized^1^. Vascular endothelial growth factor (VEGF)-A121, VEGF-A165, placental growth factor (PlGF)-1, PlGF-2, platelet-derived growth factor (PDGF)-AA, PDGF-BB, PDGF-CC, PDGF-DD, fibroblast growth factor (FGF)-1, FGF-2, FGF-6, FGF-7, FGF-9, FGF-10, FGF-18, bone morphogenetic protein (BMP)-2, BMP-3, BMP-4, BMP-7, β-nerve growth factor (NGF), neurotrophin (NT)-3, brain-derived neurotrophic factor (BDNF), insulin-like growth factor (IGF)-1, IGF-2, heparin-binding epidermal growth factor (HB-EGF), C-X-C motif ligand (CXCL)-11, and CXCL-12α were purchased from PeproTech. CXCL-12γ was purchased from R&D systems. Except for PDGF-DD and BMP-7, which were produced in eukaryotic cells, all GFs were produced in *Escherichia coli* and thus were not glycosylated. All GFs were reconstituted and stored according to the provider’s instructions to regain full activity and prevent loss of protein.

### Detection of laminin binding to recombinant GFs

ELISA tests were performed as previously reported^1^. In brief, ELISA plates (med-binding, Greiner Bio-One) were coated with 50 nM GFs at 37°C for more than 2 h. After blocking with 2% BSA solution containing PBS and 0.05% Tween 20 (PBS-T), 10 nM recombinant human laminin isoforms (−111, −211, −332, −411, −421, −511, and −521) (BioLamina) were added. Bound laminin was detected with rabbit anti-human laminin γ1 chain antibody (1:1000 dilution, Assay biotech) or rabbit anti-human laminin α3 chain antibody (1:1000 dilution, Assay biotech). After incubation with biotinylated anti-rabbit antibody for 60 min at room temperature (RT), HRP conjugated streptavidin (Jackson ImmunoResearch) was added. After 60 min of incubation at RT, 50 μL TMB substrate (Sigma-Aldrich) was added. The reactions were stopped by adding 25 μL of 2 N H_2_SO_4_. Subsequently, the absorbance at 450 nm was measured with a reference of 570 nm.

### Production and purification of recombinant laminin α3_2928-3150_ protein

Protein production and purification were performed as described previously^1^. The sequence encoding for human laminin alpha 3 LG domain Ser2928-Cys3150 (linker domain and LG4 domain) was synthesized and subcloned into the mammalian expression vector pcDNA3.1(+) by Genscript. A sequence encoding for 6 His was added at the N-terminus for further purification of the recombinant protein. Suspension-adapted HEK-293 F cells were routinely maintained in serum-free FreeStyle 293 Expression Medium (Gibco). On the day of transfection, cells were inoculated into fresh medium at a density of 1 x 10^6^ cells/mL. 1 µg/mL plasmid DNA, 2 µg/mL linear 25 kDa polyethylenimine (Polysciences), and OptiPRO SFM media (4% final concentration, Thermo Fisher) were sequentially added. The culture flask was agitated by orbital shaking at 135 rpm at 37°C in the presence of 5% CO_2_. 6 days after transfection, the cell culture medium was collected by centrifugation and filtered through a 0.22 μm filter. Culture media was loaded into a HisTrap HP 5 ml column (GE Healthcare), using an ÄKTA pure 25 (GE Healthcare). After washing of the column with wash buffer (20 mM imidazole, 20 mM NaH_2_PO_4_, 0.5 M NaCl, pH 7.4), protein was eluted with a gradient of 500 mM imidazole (in 20 mM NaH_2_PO_4_, 0.5 M NaCl, pH 7.4). The elusion solution was further purified with size exclusion chromatography using a HiLoad Superdex 200PG column (GE healthcare). All purification steps were carried out at 4°C. The expression of laminin LG domain was determined by western blotting using anti-His tag antibody (BioLegend) and the proteins were verified as >90% pure by SDS-PAGE.

### Surface plasmon resonance (SPR)

SPR analysis was performed as described previously^2^. In brief, measurements were made with a Biacore 3000 SPR system (GE Healthcare). Laminin-521 or laminin α3_2928-3150_ was immobilized via amine coupling on a C1 chip (GE Healthcare) for ~2000 or ~1000 resonance units (RU), respectively, according to the manufacturer’s instructions. VEGF-A165, PDGF-BB, or PlGF-2 was flowed at increasing concentrations in the running buffer at 20 μL/min. The sensor chip was regenerated with 50 mM NaOH for every cycle. Specific bindings of GFs to laminin were calculated by comparison to a non-functionalized channel used as a reference. Experimental results were fitted with Langmuir binding kinetics using BIAevaluation software (GE Healthcare).

### Inhibition of laminin-GF binding by heparin

ELISA plates (med-binding) were coated with 10 μg/mL laminin isoforms (−111, −211, −221, −411, −421, −511, and −521) in PBS for 2 h at 37°C. Then, wells were blocked with 2% BSA-containing PBS-T and further incubated with 1 μg/mL each of VEGF-A165, PlGF-2, or FGF-2 for 60 min at RT with 10 μM heparin. Next, the wells were incubated with biotinylated anti-VEGF, anti-PlGF, or anti-FGF-2 antibodies (R&D Systems). The antibodies were detected by streptavidin-HRP (R&D Systems). Signals were revealed and measured as described above.

### Detection of GF binding to recombinant laminin LG domain protein and the synthesized laminin HBD peptides

ELISA tests were performed as described above. In brief, ELISA plates were coated with 1 μg/mL of laminin alpha 3 LG domain recombinant protein, laminin alpha 4 LG domain recombinant protein (R&D systems), laminin alpha 5 LG domain recombinant protein (LD BioPharma), or laminin peptide (sequences are described in Table 1, chemically synthesized by Genscript) in PBS for 2 h at 37°C. 1 μg/mL of BSA served as non-binding protein control. After blocking with 2% BSA PBS-0.05% Tween 20 (PBS-T) solution, 1 μg/mL of the recombinant human proteins (VEGF-A121, VEGF-A165, PlGF-1, PlGF-2, PDGF-BB or FGF-2) or 10 μg/mL of biotinylated heparin (Sigma-Aldrich) were added. Bound GF was detected with biotinylated antibodies for human VEGF, PlGF, PDGF-BB, or FGF-2 (R&D Systems). The antibodies were detected by streptavidin-HRP (R&D Systems). Signals were revealed and measured as described above.

### Detection of recombinant syndecan binding to the synthesized laminin HBD peptides

ELISA tests were performed as described above. In brief, ELISA plates were coated with 1 μg/mL laminin peptide (sequences are described in Table 1, chemically synthesized by Genscript) in PBS for 2 h at 37°C. 1 μg/mL of BSA served as non-binding protein control. After blocking with 2% BSA PBS-T solution, 1 μg/mL of the recombinant human syndecan-1, syndecan-2, syndecan-3, syndecan-4 (all syndecan proteins are histidine-tagged; SinoBiological) were added. Bound GF was detected with anti-histidine tag antibody (1:1000 dilution, BioLegend). Signals were revealed and measured as described above.

### Cell adhesion assay

96-well plates (non-tissue culture treated, Greiner Bio-one) were pre-coated with 1 μg/mL with laminin HBD peptides in PBS for 2 h at 37°C, followed by blocking with 2% BSA PBS for 1 h at RT. Cell adhesion assays were performed using human lung fibroblasts (Lonza) in FGM-2 medium (Lonza) or human umbilical vein endothelial cells (HUVEC; Lonza) in EGM-2 medium (Lonza) supplemented with 1% fetal bovine serum (FBS) and 100 µg/mL VEGF-A165, with or without 5 mM EDTA (Sigma-Aldrich). Cells were plated at 3000 cells/well on laminin peptide pre-coated plates and incubated for 30 min at 37°C, 5% CO_2_. Then, the medium was removed, and wells were quickly washed three times with PBS. Cell numbers were quantified using a CyQUANT assay, according to the manufacturer’s instructions (Invitrogen). All cell lines were checked for mycoplasma contamination and used in passages from 5 to 8.

### Release of GF from fibrin matrix

Fibrin matrices were generated with human fibrinogen (VWF and fibronectin depleted, Enzyme Research Laboratories) as described previously^1^. In brief, fibrin matrices were generated with 8 mg/mL fibrinogen, 2 U/mL human thrombin (Sigma-Aldrich), 4 U/mL factor XIIIa (Fibrogammin; Behring), 5 mM calcium chloride (Sigma-Aldrich), 2 µM α_2_PI_1-8_-laminin peptide (sequences are described in Table 1, chemically synthesized by Genscript), and 500 ng/mL recombinant human VEGF-A165 or PDGF-BB. Thus, the peptides were incorporated into the 3D fibrin matrix through enzymatic coupling, via the coagulation transglutaminase factor XIIIa, of the α_2_PI_1-8_ peptide sequence (NQEQVSPL) fused to the laminin peptide^3–5^. Fibrin matrix was polymerized at 37°C for 1 h and transferred into 24-well Ultra Low Cluster plates (Corning) containing 500 μL of buffer (20 mM Tris-HCl, 150 mM NaCl, and 0.1% BSA; pH 7.4). A control well that served as a 100% released control contained only the GF in 500 μL of buffer. Every 24 h, buffers were removed, stored at −20°C, and replaced with fresh buffer. For the 100% released control well, 20 μL of buffer was removed each day and stored at −20°C. After 5 days, the cumulative release of GF was quantified by ELISA (DuoSet; R&D Systems), using the 100% released control as a reference.

### Mouse skin chronic wound healing model

Skin wound healing assays were performed as previously reported^1^. Briefly, C57BLKS/J-m/Lepr db (db/db) male mice were 10 to 12 weeks old at the start of the experiments. Their backs were shaved and four full-thickness punch biopsy wounds (6 mm in diameter) were created in each mouse. Directly after, fibrin matrices [80 µL total, fibrinogen (10 mg/mL), 2 U/mL human thrombin, 4 U/mL factor XIII, 5 mM calcium chloride, 2 µM α_2_PI1-_8_-α3_3_043-3067, 100 ng of VEGF-A165, and 50 ng of PDGF-BB] were polymerized on the wounds. The wounds were covered with adhesive film dressing (Hydrofilm, Hartmann). Mice were single-caged after the wound surgery. After 7 days, mice were euthanized and the skin wounds were carefully harvested for histological analysis. All animal experiments were performed with approval from the Veterinary Authority of the Institutional Animal Care and Use Committee of the University of Chicago (IACUC).

### Histomorphometric analysis of wound tissue sections

Histomorphometric analyses were performed as previously reported^1^. Briefly, an area of 8 mm in diameter, which includes the complete epithelial margins, was excised. Wounds were cut in the center into two and embedded into paraffin. Histological analysis was performed on 5 μm serial sections. Images were captured with an EVOS FL Auto microscope (Life Technologies). The extent of re-epithelialization and granulation tissue formation was measured by histomorphometric analysis of tissue sections (H&E stain) using ImageJ software. For analysis of re-epithelialization, the distance that the epithelium had traveled across the wound was measured; the muscle edges of the panniculus carnosus were used as indicator for the initial wound edges; and re-epithelialization was calculated as the percentage of the distance of edges of the panniculus carnosus muscle. For granulation tissue quantification, the area covered by a highly cellular tissue was determined.

### Statistical analysis

Statistical methods were not used to predetermine necessary sample size, but sample sizes were chosen based on estimates from pilot experiments and previously published results such that appropriate statistical tests could yield significant results. Statistically significant differences between experimental groups were determined by one-way ANOVA followed by Tukey’s HSD post hoc test with Prism software (v7, GraphPad). Variance between groups was found to be similar by the Brown-Forsythe test. For non-parametric data, the Kruskal-Wallis test followed by Dunn’s multiple comparison test was used. For ELISA data, the two-tailed Mann-Whitney U test was used. For the animal studies, experiments were not performed in a blinded fashion. Mice were randomized into treatment groups within a cage immediately before the wound surgery and treated in the same way. GF-laminin binding ELISA assays were repeated 4 times. Wound healing assays were repeated 5 times. The P values less than 0.05 are considered to be significantly different. The P values less than 0.05 and 0.01 indicate symbols * and **, respectively.

### Data availability

The data that support the findings of this study are available from the authors on reasonable request.

